# Predicting murine age across tissues and cell types using single cell transcriptome data

**DOI:** 10.1101/2022.10.19.512922

**Authors:** Janis Frederick Neumann, Ana Carolina Leote, Meike Liersch, Andreas Beyer

## Abstract

Molecular aging clocks utilize high-dimensional profiling data to predict the chronological or biological age of individuals. While this approach has proven successful across a wide range of species and tissues, the potential of using single-cell molecular profiling data for age prediction remains to be fully explored. Here, we demonstrate that aging clocks based on single-cell RNA-sequencing (scRNA-seq) data enable studying aging effects for different cell types in the same organ and for similar cell types across organs. We utilize mouse single-cell RNA-Seq data to train molecular aging clocks that distinguish between cells of young and old mice using two models: a first model trained specifically to predict the age of B cells and a second one predicting age across 70 cell types from 14 tissues.

We evaluated Elastic Net regression and two tree-based machine learning methods, Random Forest and XGBoost, as well as three distinct methods of transforming the measured gene expression values. Our models proved to be transferable to independent individuals and tissues that were not used for model training, reaching an accuracy of over 90%. A single-cell molecular aging clock trained on B cells from the spleen was capable of correctly classifying the age of almost 95% of all B cells in different organs. This finding suggests common molecular aging processes for B cells, independent of their site of residence. Further, our aging models identified several aging markers involved in translation and formation of the cytoskeleton, suggesting that these fundamental cellular processes are affected by aging independent of the cell type. Beyond showing that it is possible to train highly accurate and transferable models of aging on single-cell transcriptomics data, our work opens up the possibility of studying global as well as cell-type-specific effects of age on the molecular state of a cell.

**Graphical Abstract:** 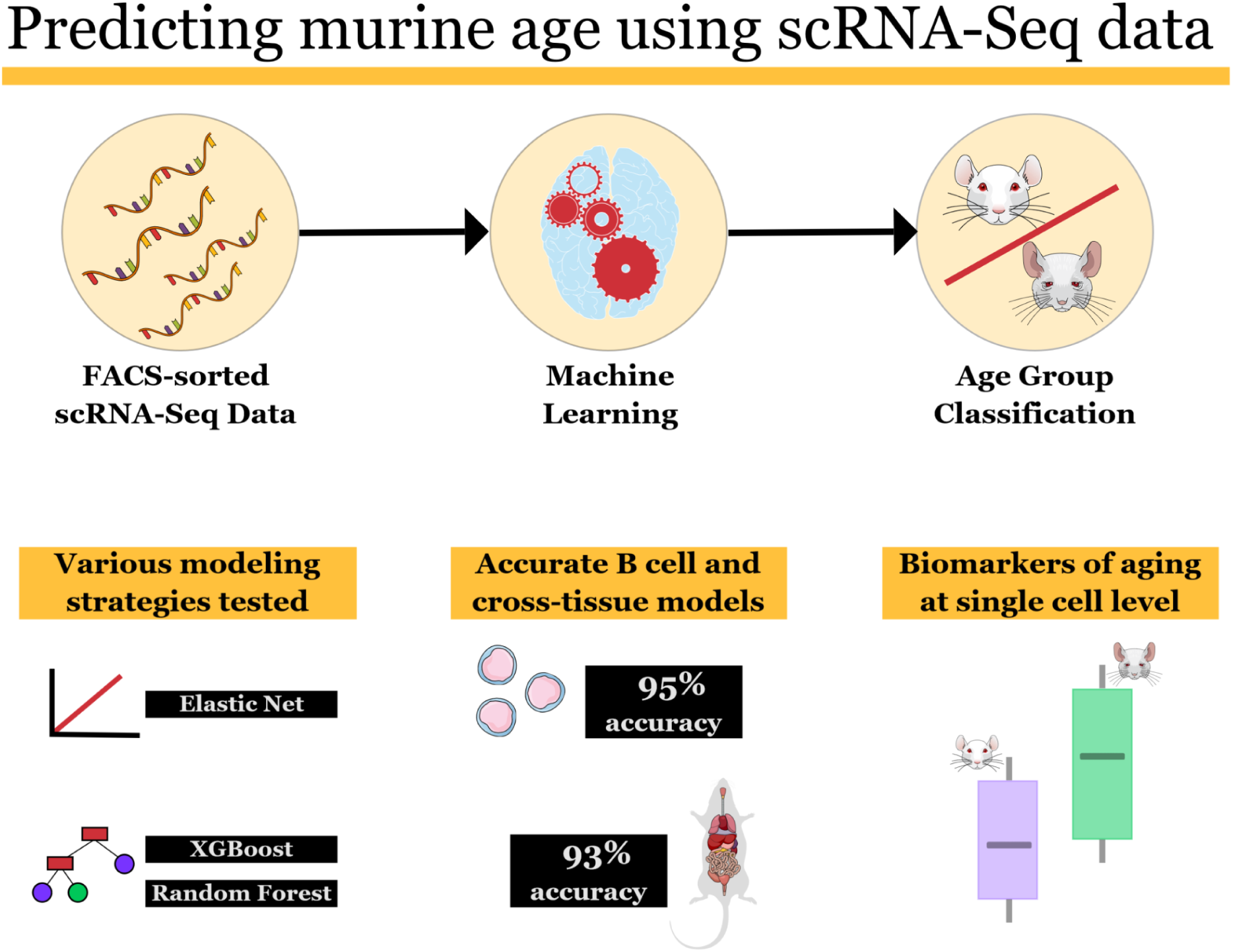

## Introduction

The last decade has seen the development and popularization of so-called ‘aging clocks’, computational models that utilize machine learning methods to predict the age of an individual (1,2). The first generation of such models were used to predict chronological age. Deviations between the age predicted by the model and the observed chronological age were interpreted as gain or loss of fitness compared to age-matched peers. Later iterations - often described as second generation clocks - were trained to model “biological” age, i. e. a measure of age-related decline using study-dependent proxy variables that are related to overall mortality (3,4). This enabled the capture of differences in aging rates between individuals and quantification of the relative deviation in biological age among individuals of the same chronological age (the time passed since birth). In fact, such deviations have been shown to be associated with mortality risk (5), supporting the use of aging clocks to guide geroprotective interventions and open new insights into the biology of aging (6). Most recently, third generation aging clocks have been developed, which measure the individual pace of aging by making use of longitudinal data to study biological age over time (7).

Aging clocks generally utilize molecular profiling data, such as transcriptomic, proteomic and metabolomic data. Using DNA methylation signatures have proven to be the most robust predictors for chronological age, enabling the application of a single aging clock across different tissues and - to certain extent - even across multiple species (8). The success of these models has led to the postulation of an “epigenetic clock theory of aging” (9–11). So far, the vast majority of aging clocks were trained on blood immune cells or bulk tissue profiling data using machine learning methods such as Elastic Net regression (12) or artificial neural networks (13). The transfer of such models between tissues has proven successful, suggesting common, generic mechanisms associated with aging across tissues (9,14). However, bulk tissue data does not provide insight into molecular aging processes in individual cell types and it remains elusive to what extent bulk aging signatures are affected by changes in cell type composition, such as immune cell invasion. Here, we show that single-cell RNA-sequencing (scRNA-seq) data has the capacity to fill this gap.

In comparison to single-cell DNA methylation data, scRNA-seq data provides two main advantages: (a) it is much more widely available and (b) it provides a simple means to distinguish cell types. While scRNA-seq data is even noisier than bulk transcriptomics data, prior work in the field has demonstrated the feasibility of the approach (15,16). However, existing work has been limited to relatively small numbers of cells and individuals or generally focused on only single cell types or tissues. It therefore remains unclear to what extent such single-cell molecular aging clocks can be used across cell types and tissues, and how the data needs to be processed for this task.

To address these questions, we made use of FACS-sorted mouse scRNA-seq data published as part of the Tabula Muris Senis data set (17). This data set comprises cells from 3, 18, 21 and 24 months old C57BL/6JN mice of both sexes, encompassing 23 tissues. We used this data to train two binary classifiers that distinguish between young (3m) and old (18m & 24m) mice. Given the age structure in the Tabula Muris Senis data set and the comparably small number of mice we decided not to predict continuous age.

The first model was trained on splenic B cells and used to predict the age of B cells in four other tissues - fat, limb muscle, liver and lung. This first model was created to investigate the molecular aging of a common cell type in different tissues, while providing specific insight into the aging immune system. Second, we trained a cross-tissue and cross-cell-type age classifier on 33,785 cells from 11 tissues and tested the model on a total of 23,675 cells, including cells taken from mice and tissues not represented in the training data. This second model enabled us to study molecular aging processes across cell-types.

In combination, these two models demonstrated the feasibility to reliably predict age using scRNA-seq data with a single, generic model. Further, our work provided insight into the aging of the same cell type in different tissues and different cell types in the same tissue, thereby identifying common molecular markers and processes affected by aging.

## Results

### B cell age prediction: predictor transformation and modeling method choice

First, we trained models distinguishing cells from young (3m) and old (18m & 24m) mice. We trained these age classifiers using Elastic Net as well as two tree-based machine learning techniques, Random Forest (18) and XGBoost (19).

Single-cell RNA-seq data suffers from multiple technical biases, requiring careful pre-processing of the data in order to make expression estimates comparable between cells and cell types. We therefore tested three pre-processing schemes using either (a) scaled quantitative expression estimates, (b) expression ranks, or (c) a binning approach grouping genes into expression quartiles, an adaptation of a binarization approach used in *C. elegans* (20).

We utilized a foldwise cross validation approach to assess how specific combinations of predictor transformation and model choice affect performance for predicting B cell age in the spleen (Fig. 1). This was done by leaving two or three mice (at least one of each of the two age groups) out of the training process for each of three folds. By performing the cross validation at the level of mice rather than cells, we could ensure that our models are not just predicting the identity of a mouse, but actually predict age, because the models had not seen any data from the left out mice during the training phase. All combinations achieved better than random predictions on average across all test folds. However, some did not do so consistently. Quartile ranking predictors proved especially susceptible to poor transferability to out-of-fold mice, falling below 8% correct predictions in the case of one particular mouse. The performance of Random Forest was slightly inferior to that of XGBoost and Elastic Net on average, achieving a mean 82.17% correct predictions compared to 88.56% achieved by XGBoost and 88.11% by Elastic Net. Random Forest also fell below 50% correct predictions (i.e. worse than random guessing) in 6 cases (out of 39) while XGBoost and Elastic Net were better than random in all but 2.

**Fig. 1.**
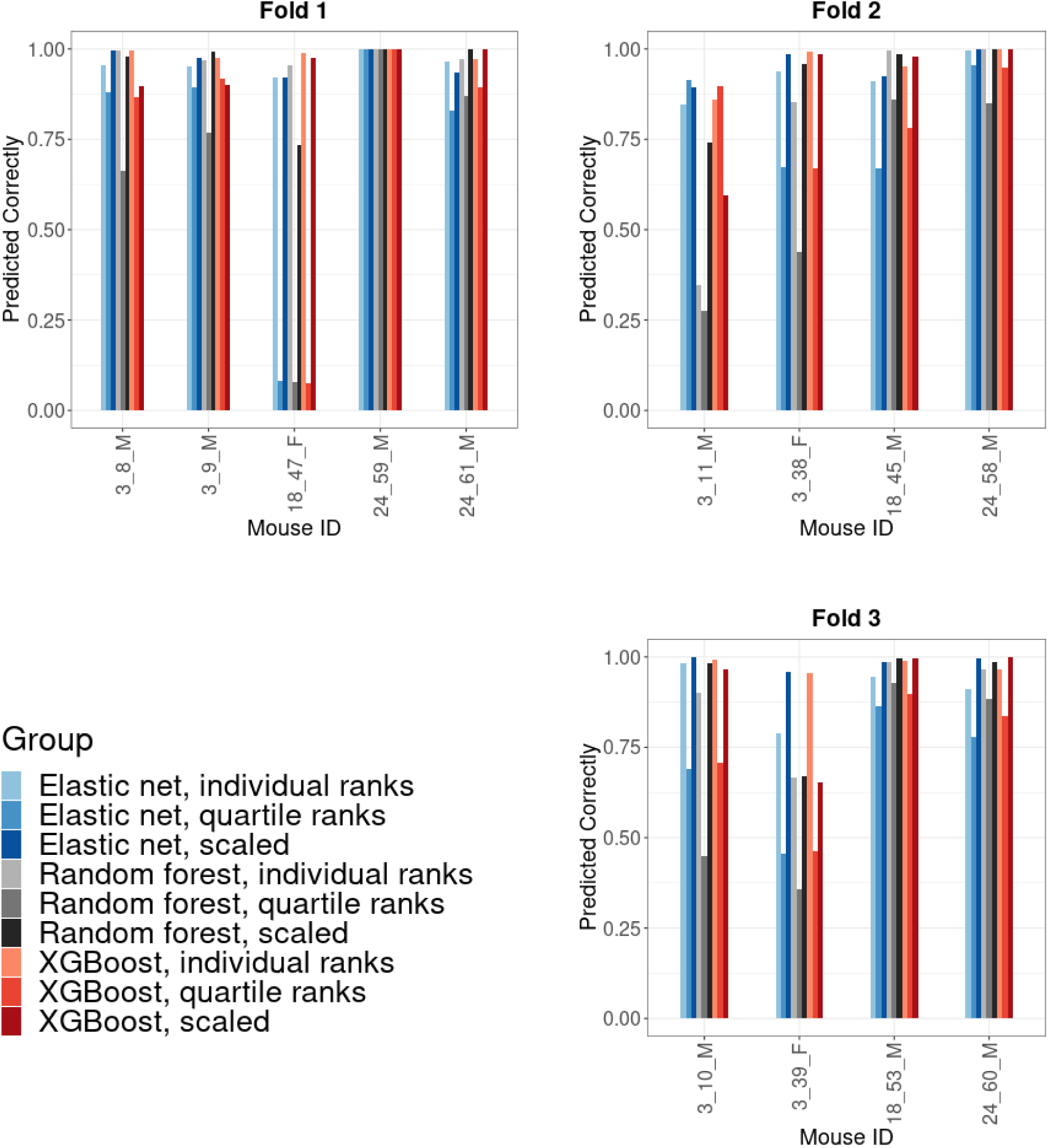
Relative performance of Elastic Net, Random Forest and XGBoost for predicting B cell age in the spleen. Each of the three subplots shows the prediction results for out-of-fold/test mice of models trained on in-fold mice of the respective fold. The x-axis labels show mouse IDs taken from the Tabula Muris Senis metadata following the pattern of {age in months}_{unique identifying number}_{sex}. Shown on the y-axis is the number of correct predictions divided by the total number of cells. Each bar represents the results for one machine learning model, with 9 models trained per fold as per the legend.

However, worse than random predictions were mostly associated with quartile rank transformation of expression values, which was used in 5 out of the 6 cases of Random Forest falling below the 50% correct predictions threshold and all cases of XGBoost and Elastic Net models doing so. The overall most successful and consistently performing strategy was using individually ranked expression values and XGBoost. This combination accomplished between 85.83% and 100% correct predictions, and 97.19% when taking the mean across mice. Elastic Net models trained on scaled predictor variables showed an almost equally good performance, with a mean accuracy of 96.70%, ranging from 89.47% to 100% correct predictions. Considering these results, we opted to focus solely on these two combinations for future analyses.

### B cell age prediction: model transfer to other tissues and age predictors

Next, we tested if scRNA-seq-based aging clocks trained on B cells from the spleen could be transferred to B cells from other tissues. For this, we used the best-performing models from above, an XGBoost model trained on individually ranked and an Elastic Net model trained on scaled expression data. Testing on left-out mice, we achieved an accuracy of 92.67% for young and 95.72% for old cells averaged across all folds in these other tissues using the XGBoost model (Fig. 2). While both prediction accuracy and the certainty of the predictions were generally stable, this did not hold up for one particular fold, where 32.24% of all young out-of-fold fat, muscle and lung B cells were misclassified as old. This can partially be traced to one mouse (ID 3_38_F) in this particular test dataset: only 50% of the cells - 50 out of 100, none of them splenic - taken from this mouse were classified correctly in this fold. In all other folds, prediction accuracy for each group was consistently above 88% irrespective of the tissue the B cells were extracted from or whether they came from out-of-fold mice.

**Fig. 2.**
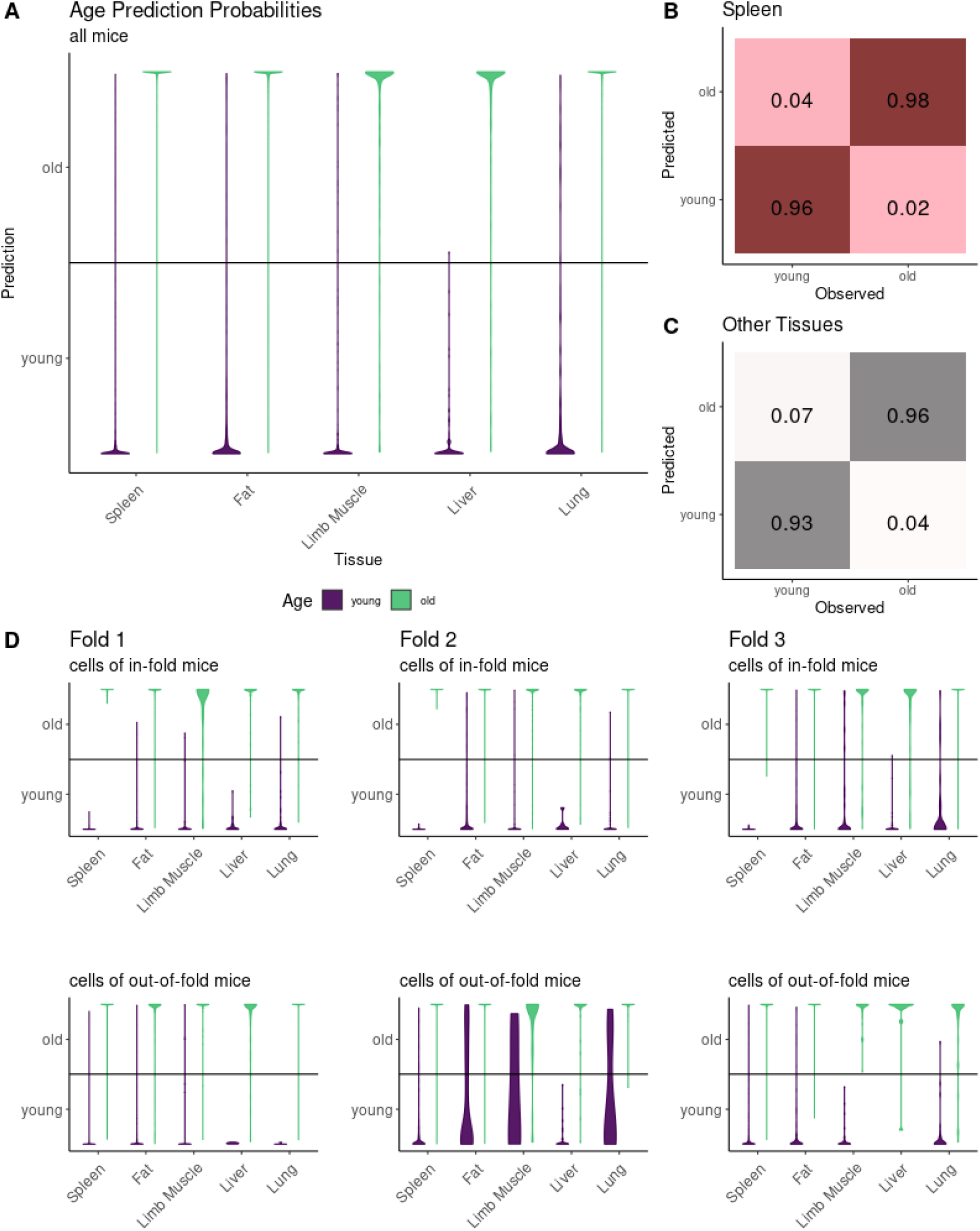
Predictive performance of the spleen-based B cell age classification model using XGBoost. Violin plots show the predictions of XGBoost on test cells by tissue and age. The y-axis shows the vote which classifies cells either as young (0/bottom) or old (1/top), with the horizontal line dividing the axis showing the demarcation between predictions (0.5). The larger violin plot shows results for all test data taken together **(A)**, while the confusion matrices show these aggregated classification results across all test cells for the spleen **(B)** as well as all other tissues combined **(C)**. The smaller violin plots **(D)** show predictions for cells from in-fold (top row) and out-of-fold mice (bottom row) separately for each fold.

The Elastic Net model performed similarly, correctly predicting the age group of 94.76% of cells from young and 94.88% of cells from old mice (Fig. 3). Similar to the XGBoost model, the poorest performance was recorded when predicting the age of cells taken from mouse 3_38_F when they were out of fold, for which an accuracy of 65% was achieved.

**Fig. 3.**
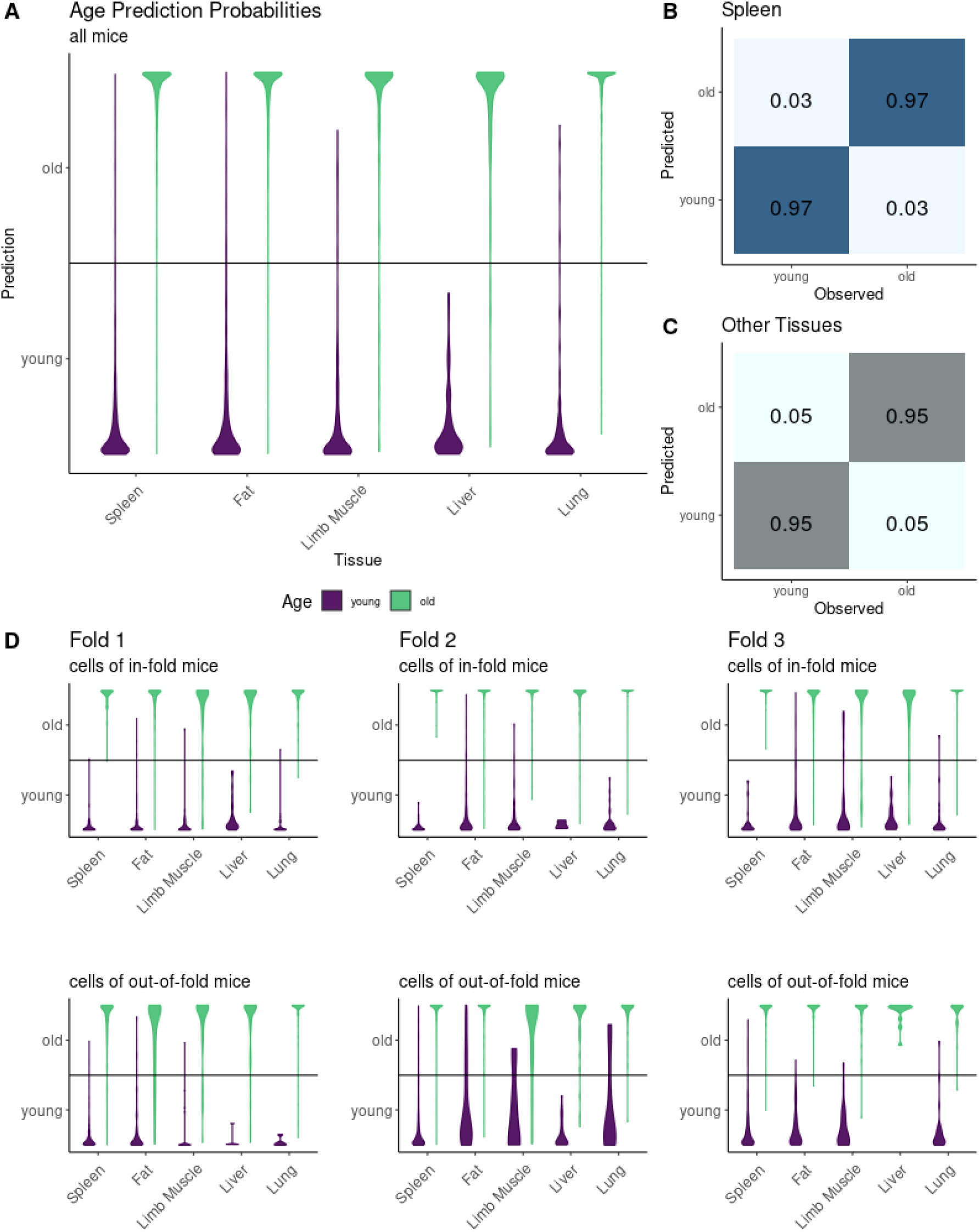
Predictive performance of the spleen-based B cell age classification model using Elastic Net regression. As before, the larger violin plot shows results for all test data taken together **(A)**, while the confusion matrices show these aggregated classification results across all test cells for the spleen **(B)** as well as all other tissues combined **(C)**. The smaller violin plots **(D)** show predictions for cells from in-fold (top row) and out-of-fold mice (bottom row) separately for each fold.

Thus, we successfully transferred scRNA-seq-based aging clocks to B cells to other tissues and the performance was only marginally worse compared to the original training tissue (spleen). These findings suggest that first, our models are not overfit to the specific training tissues and second, that general B cell aging markers seem to behave similarly independent of the tissue context.

Among the ten most important predictors learned by Elastic Net regression (Fig. 4 A) and XGBoost (Fig. 4 B) were the ribosomal genes Rpl13a, Rps28 and Rps29, the AP1 transcription factor member Jund, as well as Cfl1 and Tmsb10, which play a role in cytoskeleton organization. Among these, Rpl13a, Cfl1 and Tmsb10 were especially relevant for XGBoost, each accounting for 10% of the overall accuracy of the models trained for each fold individually. The most relevant predictor variable in the Elastic Net model was the microRNA Mir703, which has recently been found to play a role in protecting from cytotoxicity, hypoxia and inflammatory cell death (21).

**Fig. 4.**
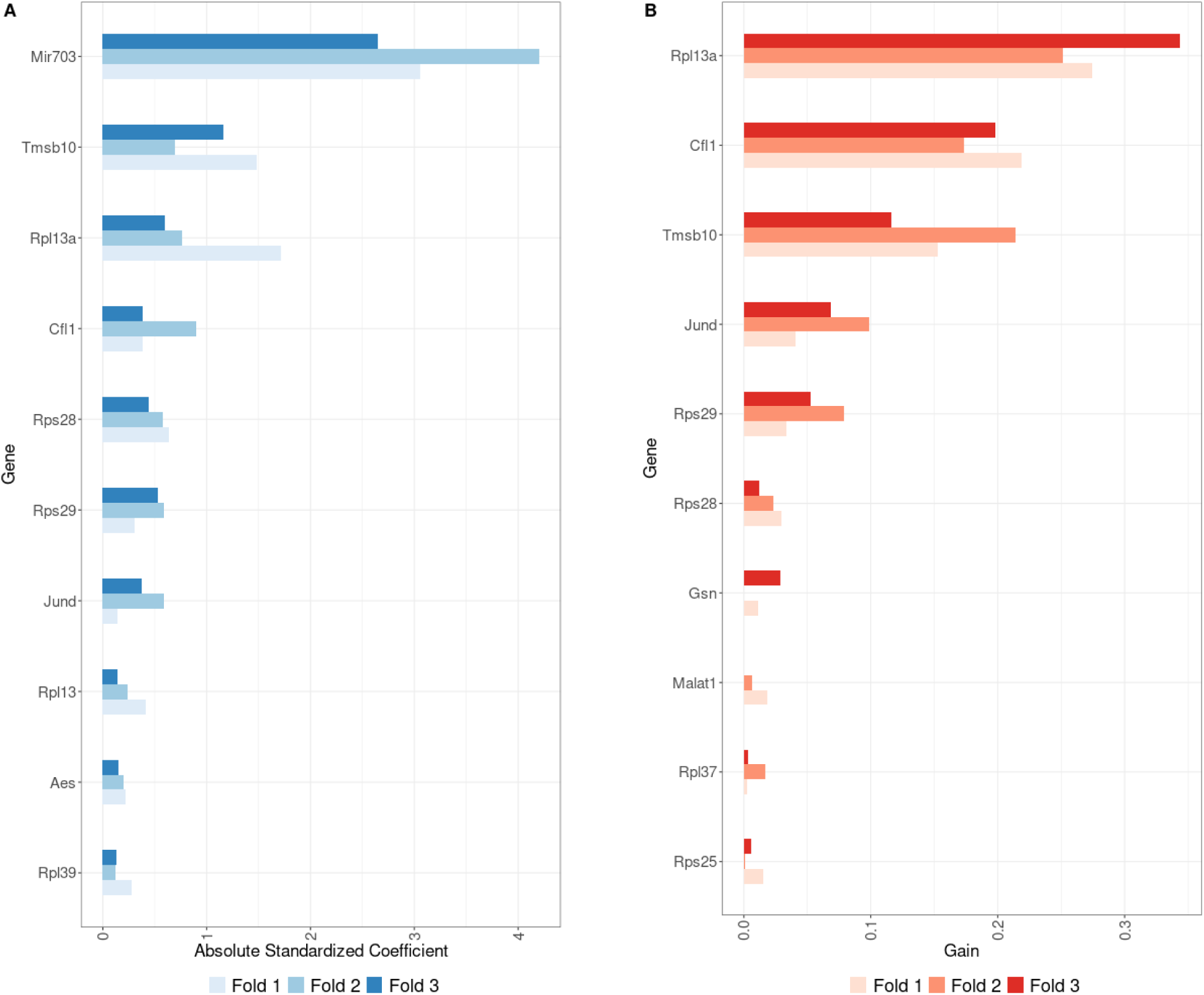
Feature importances for the most performant spleen-based B cell age classifiers using Elastic Net regression **(A)** and XGBoost **(B)**. For Elastic Net models, the absolute standardized model coefficients are used as variable importance measures. For XGBoost models, the gain associated with each feature is reported. Features are sorted by mean importance across the three models trained (one for each fold). All three Elastic Net models use the same α while all XGBoost models share the same hyperparameters.

Jund, a known proto-oncogene (22) understood to protect cells from senescence (23), was found to be upregulated in B cells of older mice (Fig. 5). Cfl1 and Tmsb10 were likewise found to be upregulated with age (Supp. Fig. 1 & 2), and Tsmb10 has been linked to senescence and apoptosis in earlier studies (24,25). Of the ribosomal genes found to be most predictive of age by both Elastic Net and XGBoost, only Rpl13a, which has also been found to be involved in the cellular inflammatory response (26), was found to be upregulated (Supp. Fig. 3). Rps28 and Rps29 were found to be downregulated with age, specifically showing a steep decline in 24 months old mice (Supp. Fig. 4 & 5). Mir703 was lowly expressed in B cells of all ages but found in higher concentrations in cells of young and almost entirely absent in older mice (Supp. Fig. 6).

**Fig. 5.**
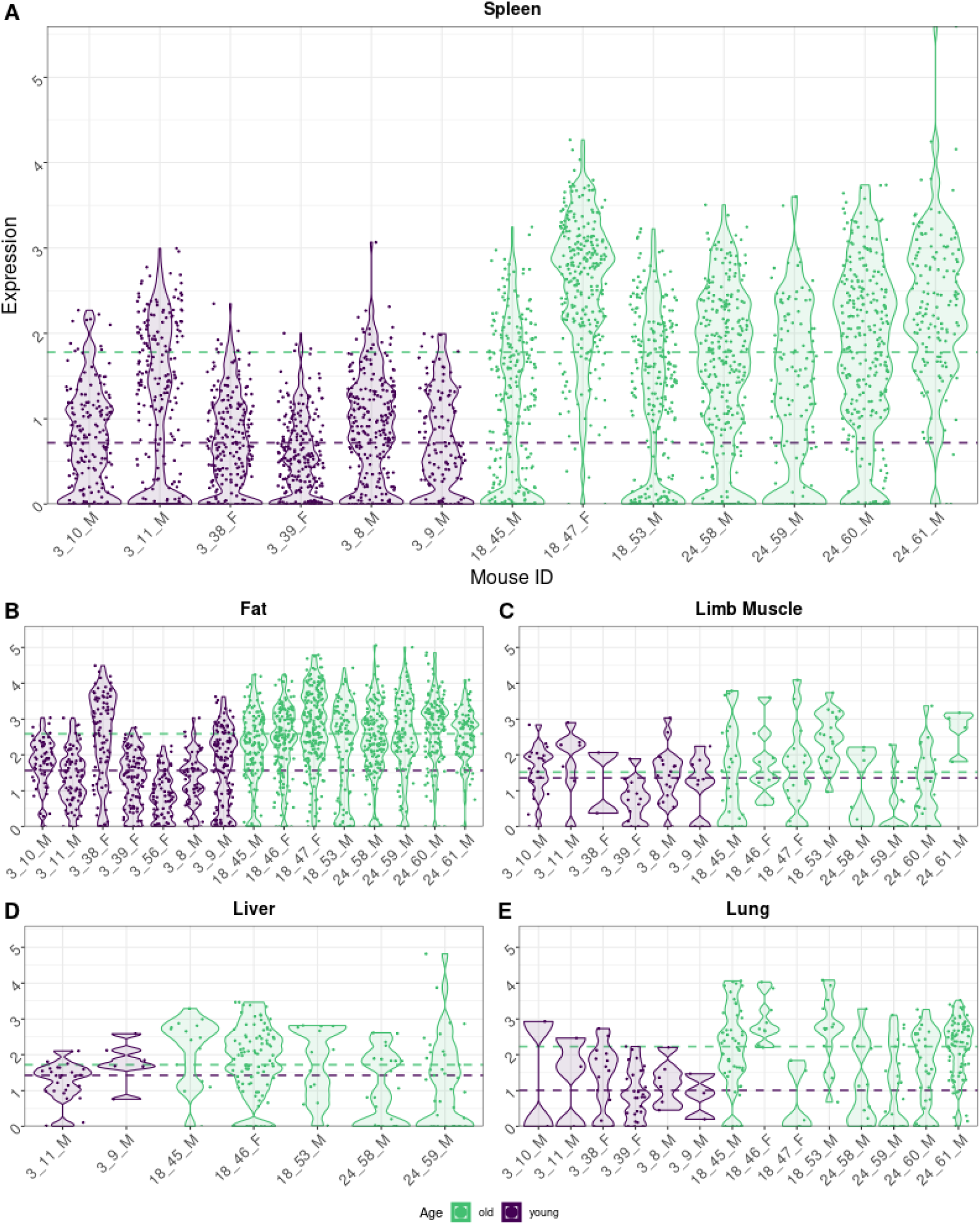
Distribution of expression values of Jund in B cells by tissue with the large panel showing expression in the spleen **(A)** and the smaller ones expression in pooled adipose tissues **(B)**, limb muscle **(C)**, liver **(D)** and lung **(E)**. The x-axis labels show mouse IDs taken from the Tabula Muris Senis metadata following the pattern of {age in months}_{unique identifying number}_{sex}. Shown on the y-axis are log10-normalized read counts. Each dot represents one cell, while violin plots show the overall distribution densities. The horizontal lines indicate median expression values by age group.

### Cross-tissue age prediction and markers

After validating the method for a single cell type, we trained XGBoost models using individually ranked gene expression values and Elastic Net models using scaled expression values on the cross-tissue data. Due to the larger data set available for this task (many more tissues and cells), we did not use a fold-wise cross validation as before. Here, we used a training set composed of cells from 15 mice - 6 young, 9 old. This data encompassed 11 tissues (bladder, brain, kidney, large intestine, limb muscle, lung, mammary gland, spleen, thymus, tongue, trachea). 10% of the cells from these mice and tissues were used as a validation set. In order to control for potential overfitting we also employed a test strategy using three further test data sets. These consisted of (a) cells from mice in the training data but three tissues (liver, pancreas, skin) that were not used during the training (i.e. ‘same mice, but different tissues’), (b) cells from two young and two old mice not used to train the model but the 11 tissues that were used for training (i.e. ‘same tissue, but different mice’), and (c) cells from the same left out mice and tissues that were not part of the training process (i.e. ‘different mice and different tissues’). Across all four test data sets taken together, the most performant XGBoost model achieved a prediction accuracy of 92.93%; 87.46% of the young cells and 96.71% of the old cells were correctly classified (Fig. 6). For test cells taken from mice and tissues that the model was trained on without the cells being used in the training data, accuracy was as high as 97.35% for young and 99.05% for old cells. The worst performance was, surprisingly, found in the test data comprising mice set aside purely for test purposes but tissues that had been used for training.

**Fig. 6.**
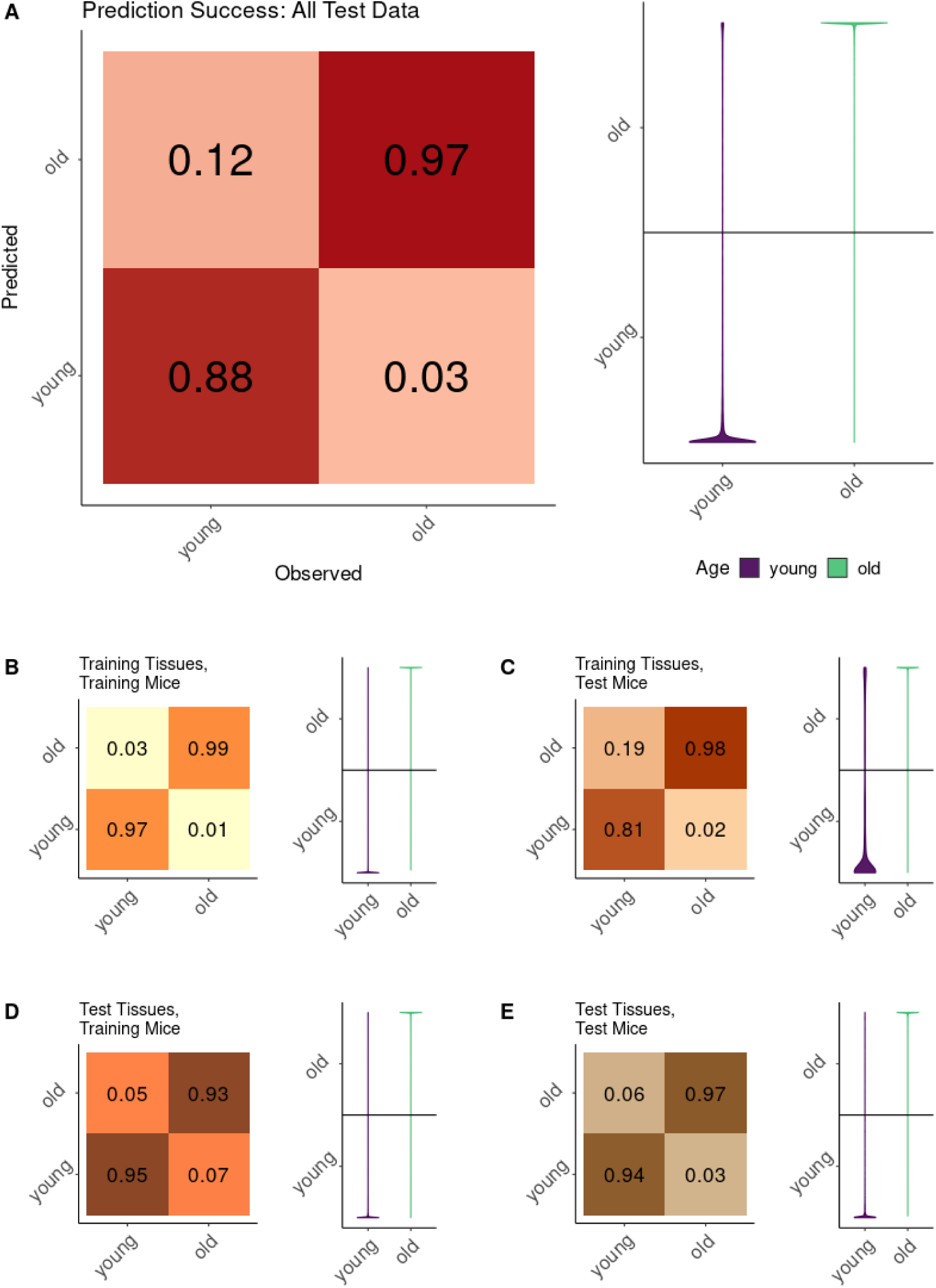
Predictive performance of the XGBoost cross-tissue age classification model for all test data taken together **(A)**, hold-out samples from tissues and mice the model was trained on **(B)**, cells from tissues the model was trained on but mice the model was not previously exposed to **(C)** or vice versa **(D)** as well as cells belonging to both mice and tissues entirely set aside for testing purposes **(E)**. Violin plots show the predictions of XGBoost by tissue and age. The y-axis shows the vote classifying cells either as young (0/bottom) or old (1/top), with the horizontal line dividing the axis showing the demarcation between predictions. Confusion matrices show the aggregated classification results.

Here, only 80.71% of the young mice in this set were predicted correctly. When tested on cells of tissues and mice not used to train the model, presumed to be the hardest samples to predict, the model’s performance was again very high: 93.68% of the young and 97.39% of the old cells in that particular test data set were predicted correctly. Since the number of mice in the test dataset is still comparably small we cannot conclude whether this unexpected finding is due to the specific nature of these individual mice or whether it is a consequence of different ‘predictabilities’ between tissues.

The Elastic Net model proved similarly successful, although its overall performance was slightly lower. It predicted 88.91% of all cells from young mice and 91.58% of all cells from old mice correctly across all four test data sets, for a total accuracy of 90.49% (Fig. 7).

**Fig. 7.**
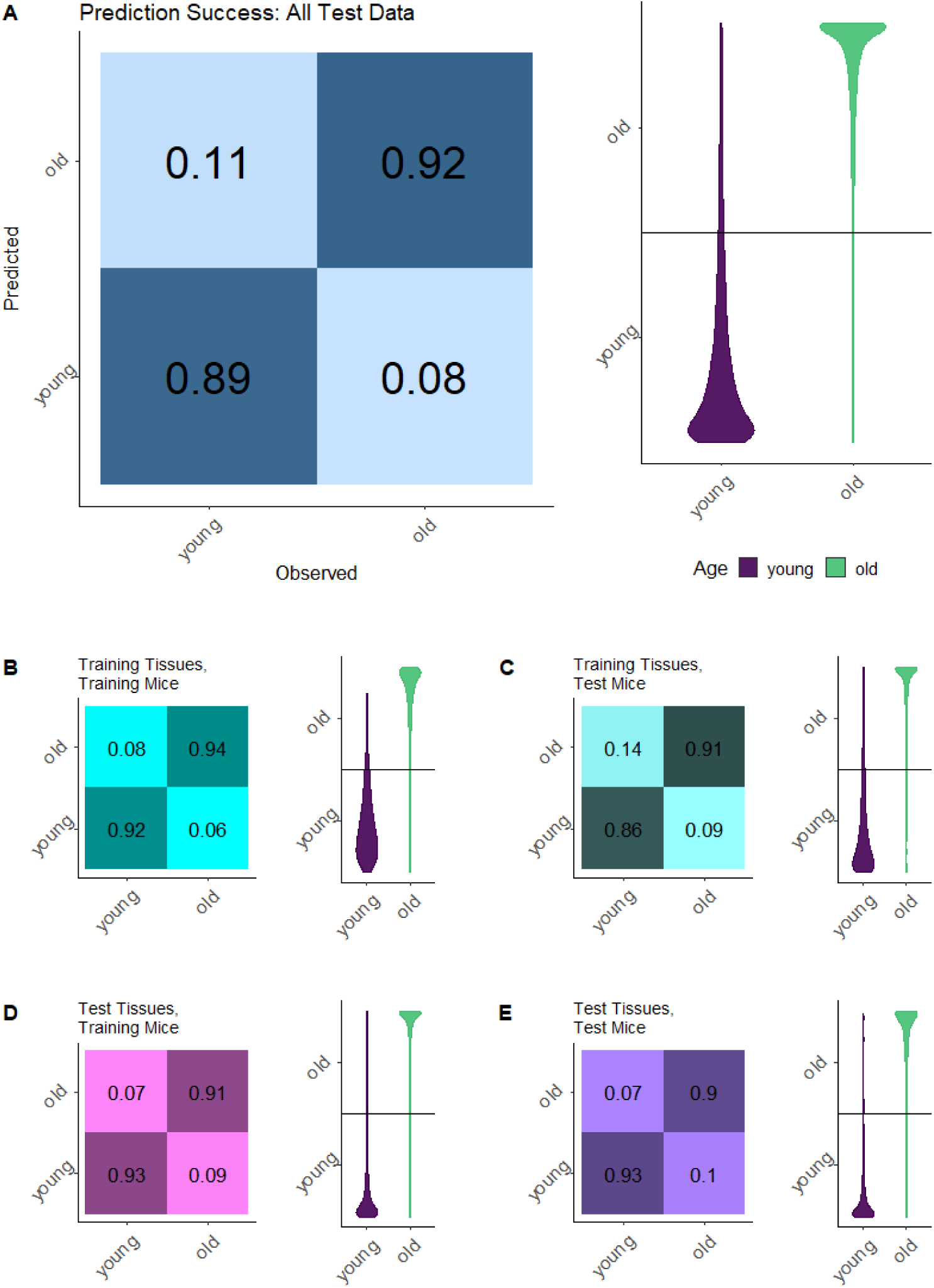
Predictive performance of the Elastic Net cross-tissue age classification model for all test data taken together **(A)**, hold-out samples from tissues and mice the model was trained on **(B)**, cells from tissues the model was trained on but mice the model was not previously exposed to **(C)** or vice versa **(D)** as well as cells belonging to both mice and tissues entirely set aside for testing purposes **(E)**.

Thus, both models showed overall a high performance, with an accuracy never dropping below 50%, i.e. always performing better than random. However, some specific combinations of tissue and age proved difficult to predict. The XGBoost model misclassified 41.32% of all lung cells from young mice (Fig. 8 A). The brain cells of young mice (all of which were microglia cells) were likewise frequently misclassified. 28.57% of them were incorrectly predicted to come from old mice. The reverse was not true; only 2.69% of the brain cells of old mice were misclassified. Strikingly, the Elastic Net model also misclassified a relatively high amount of brain cells but showed no such age bias (misclassification rate young: 15.79%, old: 14.57%; Fig. 8 B). While both XGBoost and Elastic Net struggled to classify test set samples from brain, lung and pancreas, the Elastic Net results showed less of a bias towards specific age groups here, instead wrongly classifying cells of both age groups with comparable frequency. Despite its slightly worse average performance, the Elastic Net model seemed more robust across test sets and specific tissues or ages compared to the XGBoost model (Fig. 8 B).

**Fig. 8.**
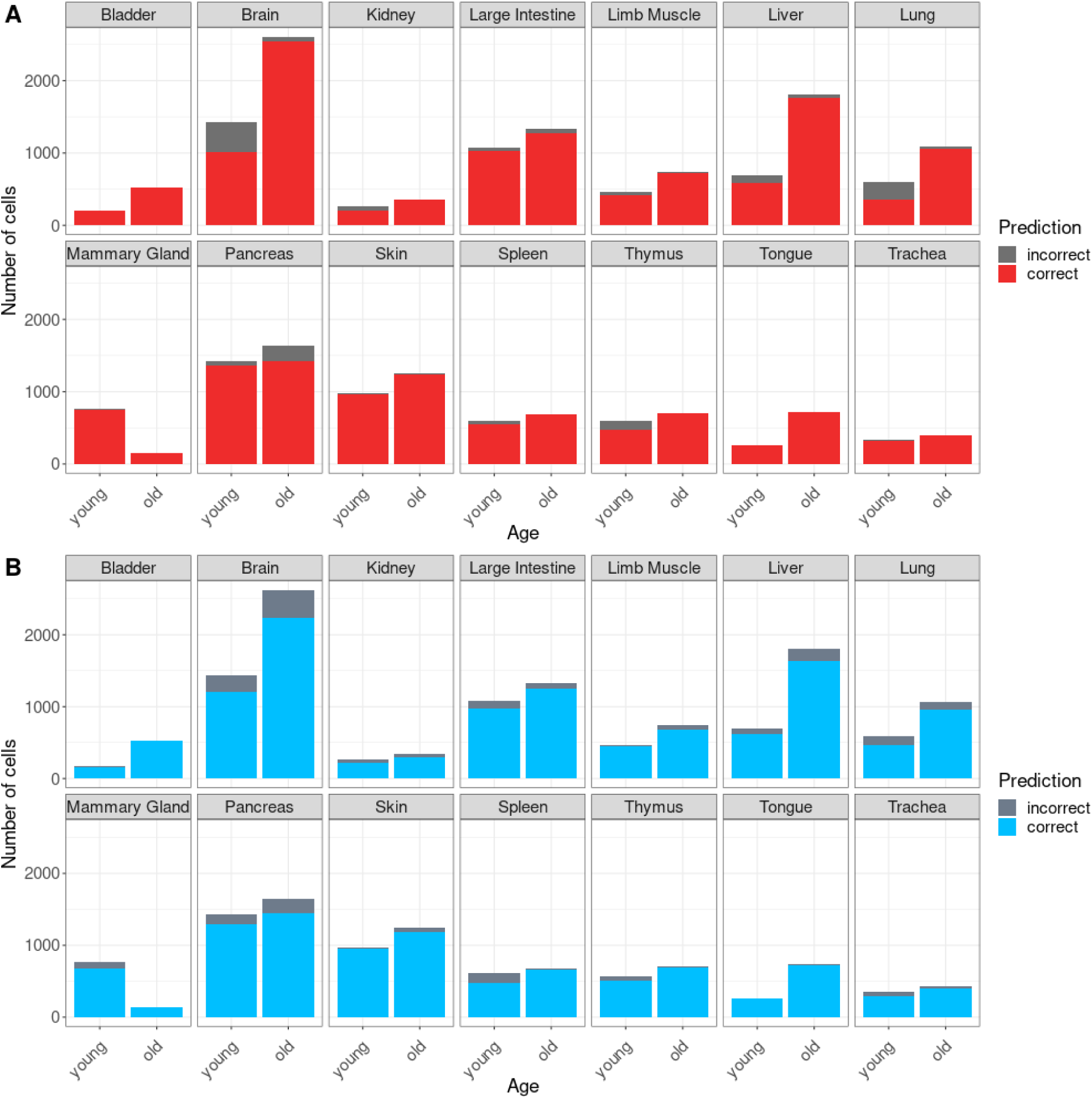
Prediction success by tissue for cross-tissue XGBoost **(A)** and Elastic Net **(B)** models. Depicted on the y-axis is the total number of cells by cell type in all four validation data sets.

The six main predictor variables shared by Elastic Net and XGBoost B cell aging models were also found to be main markers of organism aging, and their expression patterns across tissues and cell types were similar to those observed in B cells (Supp. Fig. 7). Cofilin 1 in particular was one of the three most important variables for the Elastic Net model (Fig. 9 A) and the most important one for the XGBoost model (Fig. 9 B). Tmsb10 is also the most important feature for the Elastic Net age classifier, indicating an even greater importance of biomarkers related to cytoskeletal integrity in the cross-tissue clocks. Three genes were found among the most important predictors for both cross-tissue models that did not have this level of importance for the B cell clocks. These are Aes (Tle5), a transcriptional repressor (27), the small nuclear ribonucleoprotein Snrpc and parathymosin (Ptms), all of which were found to be more highly expressed in old mice (Supp. Fig. 7). None of them have been studied extensively, however, parathymosin secretion by hypothalamic stem cells has recently been reported to protect against cellular senescence and general aging phenotypes, although the exact mechanism of action remains unclear (28).

**Fig. 9.**
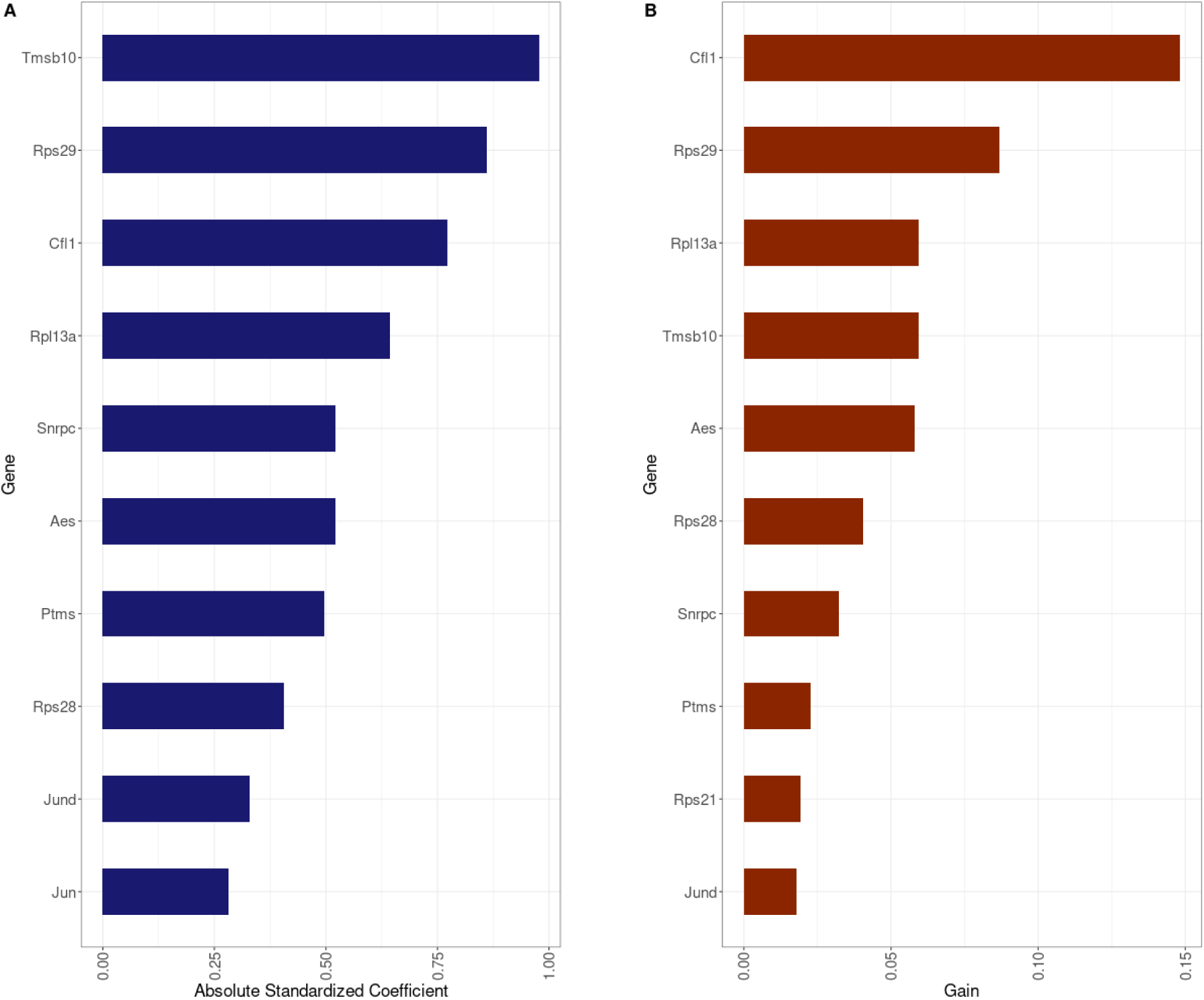
Feature importances for the most performant Elastic Net **(A)** and XGBoost **(B)** cross-tissue age classifier showing the ten most important predictor variables in order.

## Discussion

Our work demonstrates the feasibility to distinguish individual cells from old and young mice using their single-cell RNA-sequencing profiles. Moreover, the machine learning models were transferable to different mice, cell types and tissues without losing much predictive power. As we achieved an average prediction accuracy of over 90% in all test data sets in both the B cell (95.54% overall test set accuracy) and cross-tissue aging models (92.93%), our results indicate that single-cell transcriptomic aging models hold great promise for future development. While prediction accuracy fell below 90% for specific test data, these cases did not necessarily hint at an overall limitation of the approach. Rather, reduced accuracy for one set of out-of-fold mice in the B cell age classifier and cells from some tissues of mice not used for the training process in the cross-tissue model highlight a limitation of existing single-cell data sets: the number of individual organisms is limited. Differences between individuals can have a great effect on model performance by chance, as evidenced by the fact that we saw decreased accuracy for only one out of three in-fold/out-of-fold combinations of the same mice in the B cell classifier. As bigger data sets with more individuals become available, this effect should subside. However, our results do indicate that some tissues or cell types are harder to model than others or at least have an aging signature that differs from the one learned by the cross-tissue models to an extent. This can most easily be seen in microglia cells, the only brain cell type and the most abundant cell type in the data which nonetheless ranked among the most frequently misclassified by both XGBoost and Elastic Net.This difference in performance may stem from overall differences in the transcriptomic profile of microglia compared to the remaining cell types used to train our models. Another factor that may contribute to the observed performance difference is the differing quality of the scRNA-seq data available, particularly regarding factors such as dropout rate or sequencing depth.

Despite the overall high success of XGBoost, it does not outperform the more established Elastic Net regression. Neither modeling method was noticeably more successful in predicting B cell age overall, Elastic Net misclassified less samples belonging to the specific young mouse (3_38_F) that was found to be problematic to predict when out-of-fold. Similarly, the overall accuracy of the cross-tissue XGBoost model is higher than that of its Elastic Net counterpart. However, it also proved prone to biased predictions, often misclassifying young mice. While the aging models we present here proved to be highly accurate, they have several limitations. First, like most aging clocks so far, our models have only been trained and validated in one data set, although we could partially compensate for this with a sophisticated training and test data setup. Second, we could only perform basic classification differentiating between young and old mice rather than training regression models predicting the exact age of a cell. This is due to the nature of the Tabula Muris Senis FACS-sorted data, which, while offering great sequencing depth and a high amount of cells, is mainly suited to studying contrasts between young and old mice as it comprises a few discrete age groups. Third, the models we trained predict the chronological age of the mice in the TMS data. The mice were sacrificed at specific time points and there exists no data with which to infer their biological age - the physiological state as it relates to aging (10) - apart from the transcriptomic data we already used to train the classifiers. As such, it is not possible to make any definite statements about how well our aging classifiers measure, for example, age-related decline and loss of biological function. In fact, the machine learning techniques we employed naturally favored predictors that change more predictably with chronological age than other genes that undergo expression changes during the aging process but less consistently so. It is likely that the latter predictors could be better used to explain differences between the biological ages of individual cells than the more favorably selected predictors which are, by definition, used to predict chronological age. However, chronological and biological age are of course closely linked and can be expected to be correlated even where they differ (29). The now state of the art epigenetic aging clocks at first also predicted chronological age (1,9) but later evolutions of the concept were shown to be capable of modeling biological age very precisely (3).

The predictors we found can be used as biomarkers of chronological age. Among our most important predictors was the senescence marker Jund, which is a part of the AP-1 transcription factor complex. Jund-defective mice are short-lived, while Jund overexpression decreases oxidative stress and extends lifespan (30). Surprisingly, we found Jund to be consistently upregulated with age in B cells (Fig. 5), and levels of Jund are increased in older cells of most tissues in the Tabula Muris Senis data (17,31). However, it has generally been observed to be downregulated in aged organisms so far (32). Taken together, these findings suggest that the expression trajectory of a gene with age may be dependent on the cell type. For example, in the B cells upregulation of Jund may be a marker of elevated oxidative stress.

Two of the consistently most important predictors of age in our models are increased expression of Tmsb10 and Cfl1 (Supp. Fig. 1 & 2), both of which are important for cytoskeletal organization. While Tmsb10 upregulation was reported as a senescence marker (25) and accelerant of apoptosis (24), its exact molecular function is currently not understood, beyond the fact that the gene product serves as an actin binding protein. Cfl1, in contrast, is better understood.

Increased levels of Cfl1 were shown by Tsai et al. to occur in senescent cells and lead to cell cycle arrest and reduce actin depolymerization (33). Interestingly, Tsai et al. also reported increased actin polymerization rates in older cells (34), a process that is inhibited by Tmsb10, hinting at the Tmsb10 upregulation in old mice possibly being compensatory. While this is speculative, our aging models suggest that cytoskeletal organization, actin polymerization and related pathways are so far understudied vectors of cellular senescence operating across cell types.

Among the most predictive age markers found by both the B cell and cross-tissue models were transcripts encoding for ribosomal proteins. Protein synthesis undergoes a variety of changes during aging, from a decreased translation rate to aggregation of misfolded proteins (35). Loss of protein stoichiometry and a decoupling of transcript and protein levels has been identified as part of the aging process (36,37). In the Tabula Muris Senis data mRNAs of ribosomal proteins were not clearly regulated in one direction (up or down), which may in part be due to the specific functions of individual ribosomal components. L13a, for example, is not necessary for ribosomal integrity but works as a silencing regulator of translation by binding to target mRNAs (38). As part of this process, it was reported to engage the interferon-γ (IFN-γ) inflammation response, ultimately downregulating inflammatory proteins and repressing inflammatory signaling (26).

Further, Rpl13a has been reported to affect lifespan (39). We found Rpl13a to be upregulated in B cells from older mice, supporting the notion that this gene might be involved in aging-associated changes of the immune response.

It has been shown, mainly for epigenetic but also for transcriptomic data, that accurate predictions of age are possible using models fit on stochastic variation as opposed to targeted regulatory processes (40), in line with the evolutionary theory of aging of post-reproductive decline. Some of the predictors we have found show an according transcription pattern.

Tmsb10, for example, becomes increasingly abundant with age in the TMS data. However, this could simply be due to it being very lowly expressed at young age and gradually moving towards middling expression levels due to stochastic processes - essentially, noise. The increasing spread of observed values would support this. Other predictors, however, such as Rps29 and other transcripts encoding for ribosomal proteins, show increased regulation with age. This, however, could in itself be a response to accumulating damage due to prior stochastic processes. Therefore, it does not constitute proof of programmatic aging, but does show targeted regulatory changes coming with age.

In conclusion, our work demonstrates the need and the possibility to study molecular processes of aging at the level of individual cells, cell types and tissues. The single-cell molecular aging clocks presented here facilitate future research in this direction and they will serve as templates for developing new single-celll clocks on different datasets.

## Materials and methods

Raw count data was downloaded from the open data repository of the Tabula Muris Senis consortium (https://registry.opendata.aws/tabula-muris-senis/). Metadata and expression profiles were extracted from the downloaded h5ad and separately stored in feather files via *feather-format* 0.4.1 in Python 3.6.9.

### Seurat processing

Feather files containing the expression profiles of individual FACS-sorted cells for each tissue were combined with the respective metadata into a Seurat (41) object. We computed the median and median absolute deviation (MAD) of library sizes on the log2-transformed counts. Cells with library sizes above 3 MAD from the median were excluded because they potentially corresponded to doublets. Genes whose expression was not detected in more than 90% of the cells for a given tissue were excluded. After these filters, library size normalization was performed using the *LogNormalize* function of the *Seurat* R package (41) with default parameters. Rare cell types, detected only in one of the tissues and consisting of less than 100 cells passing library size filters, were removed from the analysis. The preprocessing steps detailed above were conducted on R 3.4.4 using Seurat version 3.2.0.

### Data selection

For the B cell age classifier, tissues with at least 100 B cells across all mice in the processed data were retained: Brown (BAT), Gonadal (GAT), Marrow (MAT) and Subcutaneous (SCAT) Adipose Tissue, Limb Muscle, Liver, Lung and Spleen. Due to low cell counts, all data from adipose tissues was pooled for this analysis. For the cross-tissue age classifier, all data that was retained by the Seurat processing steps was used. In both cases, the data was reduced to the subset of genes quantified and passing dropout fraction threshold filters across all considered tissues.

Retained cells were then sorted into training and test sets.

For the B cell classifiers, we employed the following cross-validation approach: Age estimators were trained on the spleen data from 6 young (3 months) and 7 old (3 18m, 4 24m) mice. We opted to use 3 folds, leaving 2 young and 2 old (1 18m, 1 24m) mice out for testing purposes in each case, with an additional 24m old mouse being used for testing in the first fold, while keeping the ratios of training to test cells and male to female mice as consistent as possible between folds (Supp. Tab. 1). Thus, every mouse was part of an independent test data set once.

For the cross-tissue model, we set aside specific mice and tissues for testing purposes, as well as retaining randomly determined specific test cells from mice and tissues generally used for training the model (Supp. Tab. 2). We thereby constructed four distinct test sets: first, cells not used for the training but belonging to mice and tissues that are part of the training data; second, cells from tissues in the training data but mice set aside for testing; third, cells from mice in training data but tissues set aside for testing; and fourth, cells from mice and tissues that were not part of the model training process. Of the 23 tissues in the Tabula Muris Senis data, we retained 14 for which we found 100 or more cells in the FACS data. Of these, three - liver, pancreas and skin - were set aside for testing purposes to ensure a diverse array of unique cell types to test on. The training data (as well as two of the validation/test sets) was made up of cells from the remaining 11 tissues: bladder, brain (microglia cells only), kidney, large intestine, limb muscle, lung, mammary gland, spleen, thymus, tongue, trachea. Of 19 total mice in the data, two old and young mice each were set aside for testing. The cells of six young and nine old mice were used for training, apart from cells from the three tissues set aside for testing purposes and 10% of the remaining cells which were set aside as validation data.

In total, 33,785 cells were retained for the training data. 3,754 cells from the same mice and cell types were set aside as a validation data set and used to optimize Elastic Net and XGBoost hyperparameters. The test data consisting of cells from the cell types the model was trained on but from left out mice comprised 12,130 cells. The test data of cells coming from mice in the training data but the three left out tissues contained 5,806 cells. The final test data set, limited to cells from mice and tissues that were not part of the training process, consisted of 1,985 cells.

### Predictor transformations

Three different predictor transformations were performed: scaling, quartile ranking and individual ranking. Scaling was performed gene-wise for each tissue separately with the *ScaleData Seurat* function (without the option to center the data being used) to constrain values to a maximum of 20 in order to reduce the potential for outliers to impact results. For individual ranks, expression ranks were computed across all genes for each cell. Ranks were assigned to genes based on the order of expression values observed in a cell, with the number of ranks equal to the number of distinct expression values. Ties were resolved by assigning the average rank. For quartile ranking, expression quartiles were computed for each cell and genes were assigned into their quartiles, meaning that the 25% lowest expressed genes attained rank 1 while the 25% of the genes with the highest expression were assigned rank 4.

### Machine learning models

For all machine learning models, the dependent variable was age, and the independent variables were all genes retained after Seurat processing.

For the B cell age classifier, 11 Elastic Net models for each fold using *α* values ranging from 0 to 1 in steps of 0.1 were trained in R 4.0.3 using *glmnet* v4.1-4 (12). For each model, the optimal *λ* value (presumed to be the *lambda.1se*, the largest *λ* within one standard error of the the *λ* resulting in the minimal mean error) were determined using 10-fold cross-validation. The optimal value of *α* was determined by calculating the accuracy for predictions on left out test cells from the same mice used to train the models (10 cells per mouse) for each fold and taking the mean accuracy. These cells were not used for any other purposes later. The optimal *α* was defined as the one for which the mean accuracy across fold was highest. This value was 0.7 for models trained on scaled expression values, 0.1 for quartile ranks, and 1 - indicating lasso regression - for individual ranks.

A similar procedure was used when training the cross-tissue model, using 33,785 cells to train the models and 3,754 cells from the same mice and cell types to select the optimal value for *α -* which was 0.9.

For all Random Forest models, age estimators consisting of 1000 decision trees were trained in R 4.0.3 using ranger v0.12.1 (42).

For the B cell and cross-tissue age classifiers, XGBoost models were trained in R 3.6.1 using *xgboost* v1.3.2.1. Optimal hyperparameters (Tab. 1) were determined using a purpose-built XGBoost framework - developed for R 3.6.1 but compatible with current R and *xgboost* versions - which can be downloaded from https://github.com/JanisNeumann/jn_xgboost.

**Table 1.**
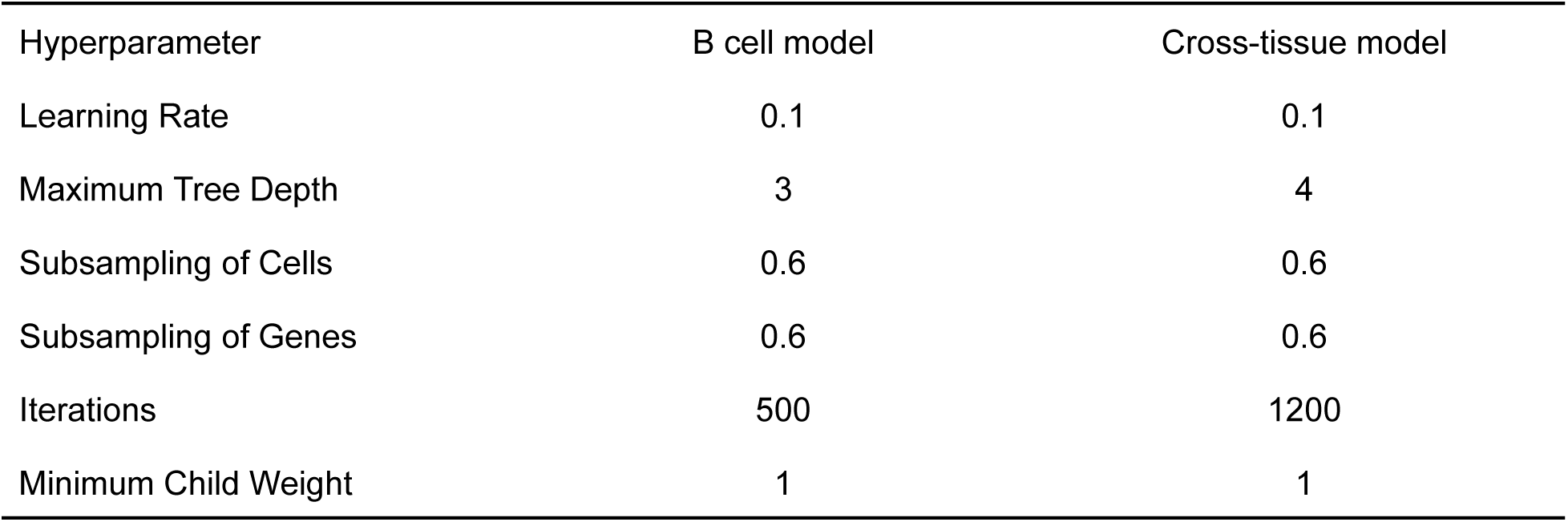
Optimal hyperparameters of B cell and cross-tissue XGBoost models.

| Hyperparameter | B cell model | Cross-tissue model |
| --- | --- | --- |
| Learning Rate | 0.1 | 0.1 |
| Maximum Tree Depth | 3 | 4 |
| Subsampling of Cells | 0.6 | 0.6 |
| Subsampling of Genes | 0.6 | 0.6 |
| Iterations | 500 | 1200 |
| Minimum Child Weight | 1 | 1 |

This framework tests all possible combinations of a user-specified array of hyperparameters. Optimal hyperparameters were found by calculating the accuracy of predictions on left out test cells as detailed for the Elastic Net models above. The hyperparameters tested for the B cell and cross-tissue age classifiers are listed in Supp. Tab. 3.

The optimal XGBoost hyperparameters were similar for the B cell-specific model and for the cross-cell-type model. A tree depth below the default of 6 used by the *xgboost* R package was found preferable in both cases. Subsampling both the cells and genes used for each tree proved successful.

### Variable Importance

To determine variable importance for the Elastic Net models, the standardized model coefficients were used. Standardization was achieved by multiplying the coefficient associated with each predictor variable with the standard deviation of that variable in the training data (43). The absolute values of these standardized coefficients are reported as the final variable importance.

The variable importance measure used to evaluate potential aging markers found by XGBoost is Gain. Gain is inherently a split-based metric, with the Gain of each individual split being defined as

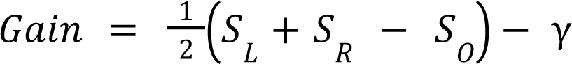

Here, S_L_ and S_R_ are the scores of the new left and right leaf after the split, while S_O_ is the score of the original leaf. The score is dependent on the particular loss function used. Regularization parameter γ is an optional way of penalizing the generation of additional splits, enforcing a minimal loss reduction. Our models utilized “multi:softprob” classification. The most successful models trained made no use of γ.

In terms of the full XGBoost model, the Gain associated with a predictor is its relative contribution to the prediction of the training data. To arrive at this number, all individual Gain values of splits using the predictor are added up and divided by the total number of splits, leading to a value between 0 and 1 - 0 indicating an unused predictor while 1 is practically impossible to achieve.

## Acknowledgments

JN received funding from the Deutsche Forschungsgemeinschaft (DFG), German Research Foundation, (CRC 1310). ACL received support by the Cologne Graduate School of Ageing Research, funded by the Deutsche Forschungsgemeinschaft (DFG), German Research Foundation, under Germany’s Excellence Strategy - EXC 2030/1 - 390661388.

