## Supplemental Figures for "Predicting murine age across tissues and cell types using single cell transcriptome data"

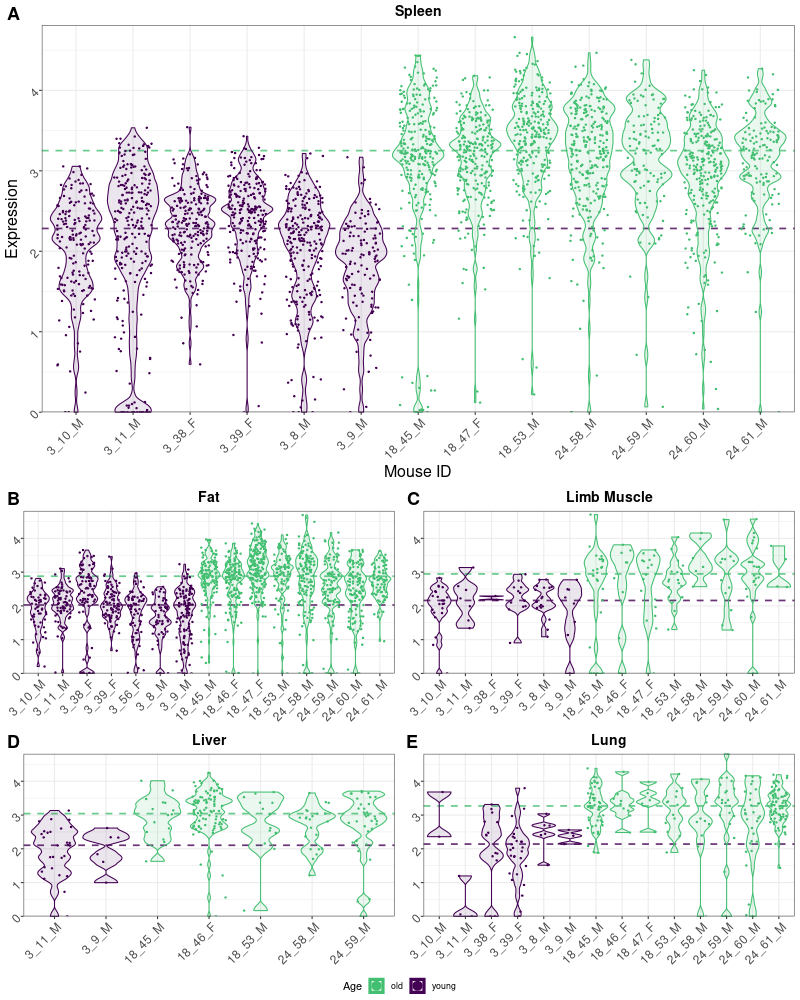

**Supp. Fig. 1.** Distribution of expression values of Cfl1 in B-cells by tissue with the large panel showing expression in the spleen **(A)** and the smaller ones expression in pooled adipose tissues **(B)**, limb muscle **(C)**, liver **(D)** and lung **(E)**. The x-axis labels show mouse IDs taken from the Tabula Muris Senis metadata following the pattern of {age in months}_{unique identifying number}_{sex}. Shown on the y-axis are log10-normalized read counts. Each dot represents one cell, while violin plots show the overall distribution densities. The horizontal lines indicate median expression values by age group.

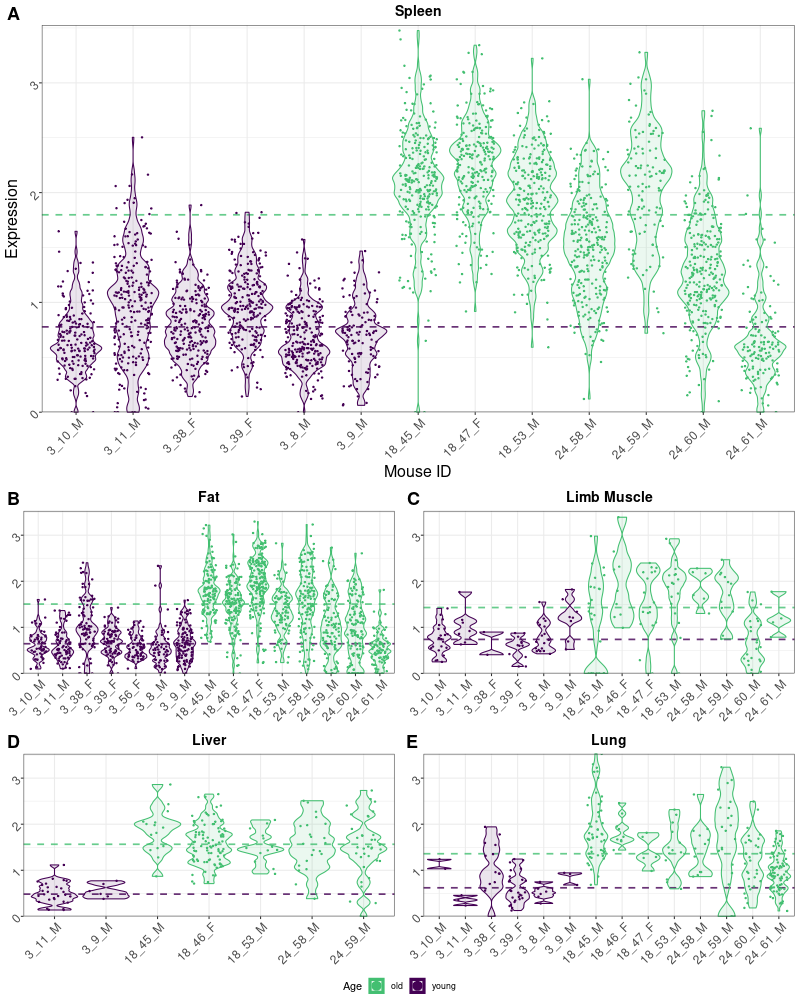

**Supp. Fig. 2.** Distribution of expression values of Tmsb10 in B-cells by tissue.

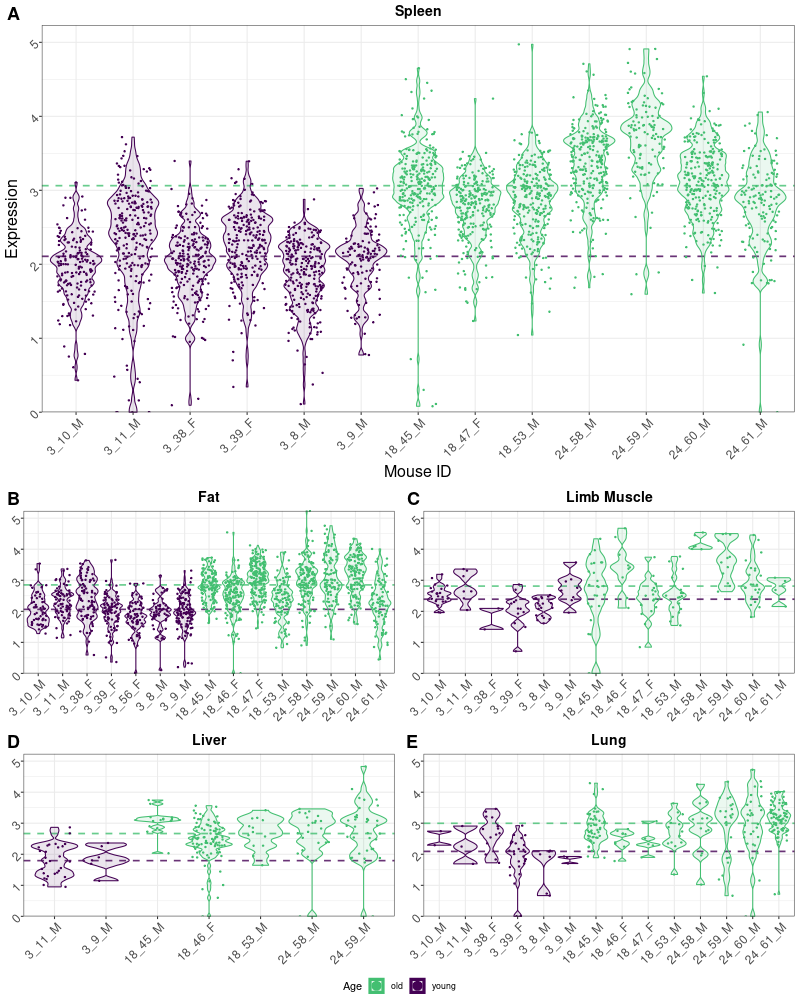

**Supp. Fig. 3.** Distribution of expression values of Rpl13a in B-cells by tissue.

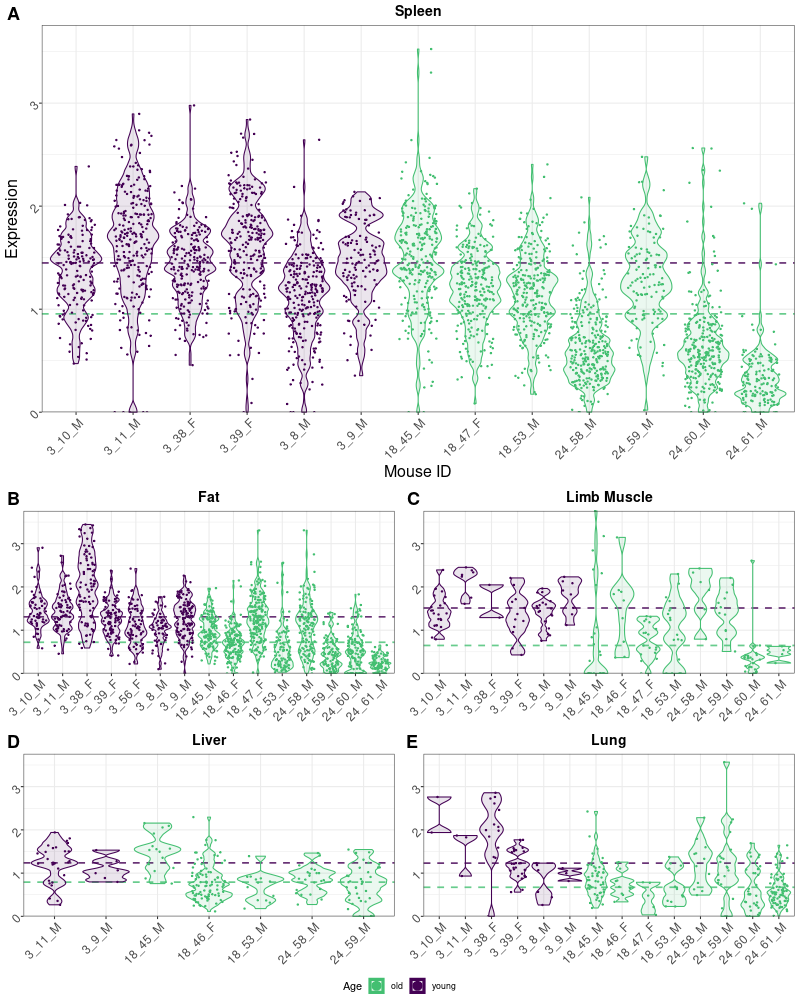

**Supp. Fig. 4.** Distribution of expression values of Rps28 in B-cells by tissue.

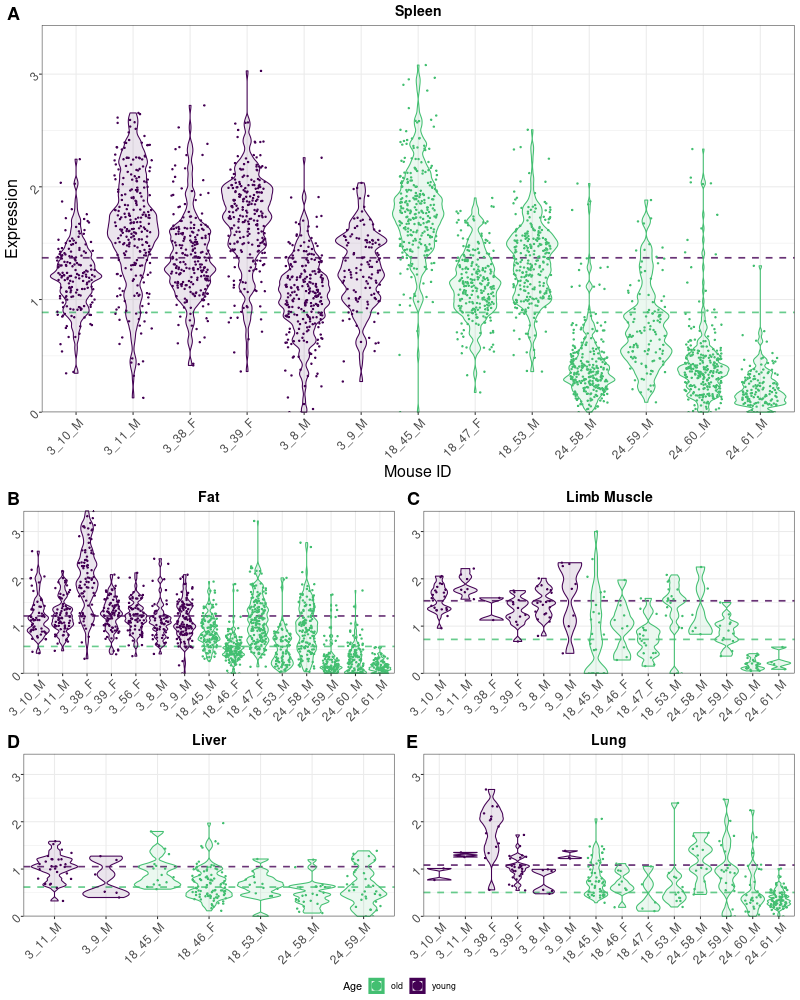

**Supp. Fig. 5.** Distribution of expression values of Rps29 in B-cells by tissue.

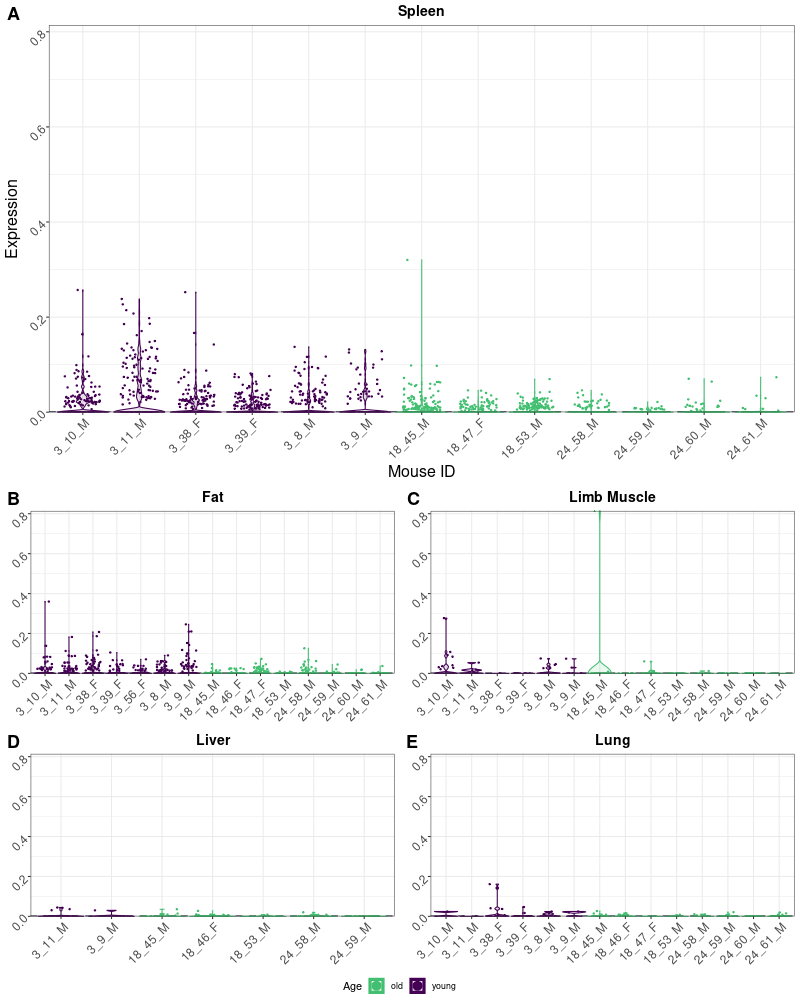

**Supp. Fig 6.** Distribution of expression values of Mir703 in B-cells by tissue.

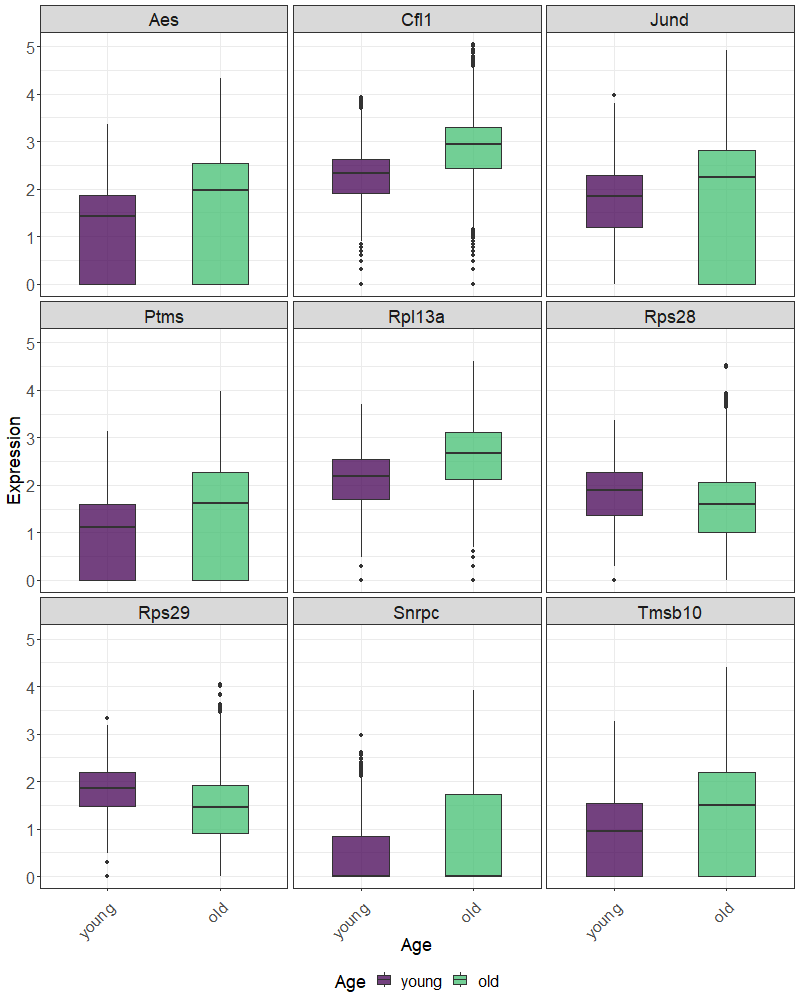

**Supp. Fig. 7.** Distribution of expression values of all genes found among the ten most important predictor variables in both the XGBoost and elastic net cross-tissue model. The x-axis labels show age. Shown on the y-axis are log10-normalized read counts.
